# Quantifying Cerebellar Signal Detectability in MEG and EEG in Epilepsy Using Anatomically Informed Source Modeling

**DOI:** 10.64898/2026.01.14.699512

**Authors:** Teppei Matsubara, Abbass Sohrabpur, Seppo Ahlfors, Mainak Jas, John Samuelsson, Padmavathi Sundaram, Steven Stufflebeam

## Abstract

**Objective:** The cerebellum is increasingly recognized as a key component of large-scale brain networks implicated in epilepsy, yet its electrophysiological characterization remains limited in noninvasive recordings. This limitation arises from the cerebellum’s depth, complex folding, and unfavorable source orientations, which challenge conventional magnetoencephalography (MEG) and electroencephalography (EEG). Here, we quantitatively characterize cerebellar signal detectability across modalities and sensor configurations using anatomically informed source modeling at the population level.

**Methods:** We analyzed clinical MEG and EEG recordings from a large cohort of patients with epilepsy undergoing presurgical evaluation. Cerebellar and cerebral source spaces were constructed using subject-specific anatomical models derived from routine clinical MRI, enabling consistent forward modeling across individuals. Signal-to-noise ratio (SNR) was estimated at individual source locations and summarized at the regional level. In addition to clinical Superconducting quantum interference device (SQUID)-MEG and EEG, multiple on-scalp optically pumped magnetometer (OPM) configurations were evaluated through simulation, including layouts matched to clinical sensor geometries and layouts optimized for posterior fossa coverage. The effects of source orientation, sensor-source distance, and head size on SNR were systematically investigated.

**Results:** In routine clinical recordings, cerebellar SNR was consistently lower than superficial cortical reference levels, confirming the limited detectability of cerebellar activity with standard SQUID-MEG and EEG. Reducing sensor-source distance by placing OPMs at SQUID-equivalent locations, i.e., projecting SQUID sensor locations to the scalp, did not improve cerebellar SNR, indicating that proximity alone is insufficient for better detectability of deeper sources. In contrast, cerebellar-optimized OPM layouts produced substantial SNR gains in posterior cerebellar regions. The effects of source orientation influence SNR differences between OPM and EEG (under identical sensor/electrode coverage) but were secondary to depth- and geometry-related constraints. Mediation analysis further demonstrated that relative sensor distance significantly mediated OPM-related advantages in posterior cerebellar regions, particularly in individuals with smaller head sizes.

**Conclusions:** These findings demonstrate that cerebellar signal detectability is governed primarily by anatomical depth and geometry rather than sensor proximity alone. Anatomically informed source modeling, combined with flexible and region-specific sensor layouts, enables meaningful improvements in cerebellar SNR that are not achievable with fixed-helmet systems. While directly motivated by epilepsy, this framework advances human brain mapping beyond the cerebrum by providing a principled approach for evaluating MEG and EEG sensitivity in deep and highly folded brain structures.

## 1. Introduction

Epilepsy is increasingly recognized as a disorder of large-scale neural networks that extend beyond cerebral cortex (Spencer, 2002). Among these distributed structures, the cerebellum has attracted growing attention for its potential involvement in seizure-related networks (Elder, Kerestes, Opal, Marchese, & Devinsky, 2025; Streng, 2023; Streng et al., 2025; Streng & Krook-Magnuson, 2021). Although traditionally viewed as a center for motor coordination, the cerebellum maintains extensive reciprocal connections with cortical and subcortical regions implicated in epilepsy (Kandel & Buzsaki, 1993; Krook-Magnuson, Szabo, Armstrong, Oijala, & Soltesz, 2014; Kros et al., 2017; Mitra & Snider, 1975), suggesting a possible role in seizure propagation, modulation, and disease progression (Blumenfeld et al., 2009; Caveness, Kosaka, Hosokawa, & O’Neill, 1977; Kandel & Buzsaki, 1993; Kros et al., 2017; Mitra & Snider, 1975; Niedermeyer & Uematsu, 1974). Importantly, accumulating neuroimaging evidence indicates that cerebellar involvement in epilepsy is region-specific (Hagemann et al., 2002; Oyegbile et al., 2011; Sone, Sato, Shigemoto, Kimura, & Matsuda, 2022). A large-scale structural MRI study of patients with epilepsy showed that cerebellar volume loss is most pronounced in posterior, predominantly non-motor lobules, such as Crus I/II and lobule VIIB, while the anterior sensorimotor lobe, vermis, and flocculonodular lobe are relatively spared across epilepsy syndromes (Kerestes et al., 2024). These posterior cerebellar regions are strongly connected with association and limbic cortices and have been implicated in cognitive and behavioral comorbidities commonly observed in patients with epilepsy (Elder et al., 2025). Despite this emerging perspective, cerebellar electrophysiology in humans remains poorly characterized.

Investigating cerebellar activity using noninvasive electrophysiology is technically challenging. The cerebellar cortex lies deep within the posterior fossa and exhibits an exceptionally folded geometry, complicating forward modelling and reducing sensor sensitivity by signal cancellation effects and large source-sensor distances (J. G. Samuelsson, Sundaram, Khan, Sereno, & Hämäläinen, 2020). Routine presurgical evaluations using electroencephalography (EEG) and Superconducting quantum interference device (SQUID)-based magnetoencephalography (MEG) are primarily optimized for cerebral coverage. Even clinical magnetic resonance imaging (MRI) may incompletely cover inferior cerebellar regions, limiting accurate anatomical modeling and coregistration.

Signal-to-noise ratio (SNR) is a central determinant of detectability in noninvasive electrophysiological recordings(J. G. Samuelsson, Peled, Mamashli, Ahveninen, & Hamalainen, 2021). Previous studies comparing SNR between SQUID-MEG and EEG have largely focused on the cerebrum and have typically enrolled only small numbers of subjects (Ahlfors, Han, Belliveau, & Hämäläinen, 2010; Goldenholz et al., 2009; Hill et al., 2020; Hunold, Funke, Eichardt, Stenroos, & Haueisen, 2016; Iivanainen, Stenroos, & Parkkonen, 2017; Marhl, Jodko-Wladzinska, Bruhl, Sander, & Jazbinsek, 2022; Piastra et al., 2021). Such limited samples cannot adequately capture interindividual variability in head size, anatomy, and sensor positioning, all of which strongly influence measured signals. In particular, source orientation plays a critical role in determining relative sensitivity between MEG and EEG (Hämäläinen, Hari, Ilmoniemi, Knuutila, & Lounasmaa, 1993), whereas source depth is a dominant factor when comparing on-scalp optically pumped magnetometer (OPM) configurations with conventional SQUID systems. Population-level estimation of SNR across a large clinical cohort, in contrast, provides a more reliable and generalizable understanding of modality performance under realistic clinical conditions.

This population-based perspective is especially important for the cerebellum. Cerebellar geometry and source orientation can vary markedly across individuals, and the depth of cerebellar sources places them at the limits of detectability for conventional systems. To date, no systematic, large-scale evaluation of cerebellar SNR across electrophysiological modalities has been established, limiting both the mechanistic understanding of cerebellar signal propagation and measurement at sensor level (based on biophysical principles) as well as its translation to clinical practice.

Recent advances in sensor technology and anatomical modeling provide an opportunity to address this gap. OPMs can be positioned directly on the scalp, eliminating the constrains of traditional cryogenic SQUID helmets and reducing the sensor-source distance (Alem et al., 2023; Bagic et al., 2023; Boto et al., 2018; Jas, Jones, & Hämäläinen, 2021; Pedersen, Abbott, & Jackson, 2022; Sheng et al., 2017). This flexibility enables individualized sensor layouts that may better target deep structures such as the cerebellum, raising the possibility of improved detectability compared with fixed-geometry systems. However, whether proximity alone is sufficient to overcome depth-related limitations remains unclear.

In parallel, advances in automated neuroanatomical modeling now allow more accurate reconstruction of the cerebellum from routine clinical MRI. Our framework, ARCUS (John G Samuelsson, Rosen, & Hämäläinen, 2020), was developed using a high-resolution ex vivo cerebellar template to preserve the folding geometry that is otherwise impossible to resolve in standard MRI (Sereno et al., 2020). By diffeomorphically registering this template to individual anatomy, ARCUS provides an approximate but highly detailed cerebellar source space that captures the cerebellar cortical folding, making it suitable for electrophysiological forward modeling. When combined with realistic forward and noise modeling, this approach enables systematic estimation of modality-specific SNR in a matter that reflects each patients’ anatomy.

In this study, we apply ARCUS to clinical MRI data from patients with epilepsy to evaluate cerebellar SNR across EEG, SQUID-MEG and simulated OPM-MEG configurations. Cerebral SNR is quantified at the population level to provide a reference for overall modality performance, whereas detailed region-based analyses are focused on the cerebellum. Using simulation-based forward modeling, we assess how sensor modality, geometry, coverage, source orientation, and depth jointly influence detectability, and whether cerebellar-optimized OPM layouts can improve sensitivity relative to conventional systems.

By focusing on modality-specific and region-specific cerebellar SNR and its anatomical determinants, this work establishes a framework for individualized cerebellar electrophysiology in the patients with epilepsy. More broadly, it demonstrates how anatomically informed modeling combined with flexible sensor technology extends noninvasive electrophysiological sensitivity beyond the cerebrum, toward measuring deep brain structures that have traditionally remained difficult to assess in clinical practice.

## 2. Materials and Methods

### 2.1 Subjects

This retrospective study included patient with intractable epilepsy who underwent routine presurgical evaluation with clinical SQUID-MEG between May 2021 and May 2022 at the Athinoula A. Martinos Center for Biomedical Imaging (Matsubara et al., 2024). All recordings were acquired as part of standard clinical care. This study is approved by Massachusetts General Brigham’s institutional review board (#2010P001169).

### 2.2 Data Acquisition

MEG recordings were obtained using standardized clinical procedure (Tanaka, Ahlfors, & Stufflebeam, 2024). Participant were positioned supine inside a three-layer magnetically shielded room, with a head support spoon and additional cushioning to maximize comfort and minimize the head motion. To optimize signal quality, the head was carefully positioned at the center of the MEG helmet and seated as deeply as possible within the rigid helmet. Continuous recordings of approximately 50 minutes were acquired.

When tolerated, a 70-channel EEG cap (EasyCap) was applied for simultaneous EEG recording, together with electrooculogram and electrocardiogram. Some patients declined EEG measurement.

Recordings were performed with a whole-head 306-channel cryogenic SQUID system (VectorView, Elekta Neuromag, Helsinki, Finland), comprising 102 magnetometers and 204 planar gradiometers. Polhemous digitization was used to record EEG electrode locations and additional scalp surface points. For interictal spike localization and anatomical modeling, individual T1-weighted MRI scans acquired at referring institutions were used. Because MRI data were obtained clinically across multiple sites, acquisition parameters and image quality varied.

For the cerebellar reconstruction (Section 2.4), individual MRI scans were first processed with Freesurfer (version 7.4.1, standard parameters). Because ARCUS relies on anatomically valid whole-brain surface reconstruction as a prerequisite, initial inclusion and exclusion criteria were defined to ensure sufficient image quality for standard Freesurfer processing at the cerebral level. Inclusion criteria comprised of; (i) availability of individual T1-weighted MRI; and (ii) isotropic or near-isotropic voxel size < 5 mm. A total of 109 patients initially met these criteria. Exclusion criteria included; (i) gross cerebral or cerebellar abnormalities (e.g., tumor, surgical cavity, cyst); (ii) failure of FreeSurfer reconstruction due to artifacts (e.g., shunt) or poor image quality insufficient for reliable whole-brain surface reconstruction required for clinical MEG analysis; (iii) inability to construct a valid three-layer boundary element (BEM) model due to incomplete head coverage; or (iv) inability to acquire EEG.

After applying these criteria, 62 patients were included for the subsequent ARCUS processing (mean ± SD age = 26.2 ± 14.0 years, range 4.7–61.4 years; 25 males). MRI data were acquired on 3T scanners in 59 patients and on 1.5 T scanners in 3 patients. Slice thickness was < 1 mm in 19 patients, 1–1.2 mm in 31 patients, and ≥ 1.2 mm in 12 patients (range 0.84–4.5 mm). Among excluded patients, exclusion reasons comprised of structural abnormalities (n = 9), FreeSurfer’s failure to reconstruct surfaces or poor image quality (n = 18), BEM construction failure (n= 11), and inability to acquire EEG (n = 10).

### 2.3 Forward modeling

The MEG and EEG forward models were constructed based on neural current sources located on cortical surfaces. In the cerebrum, dendritic current in pyramidal neurons are modeled as source current dipoles, oriented perpendicularly to the cortical mantle (Gutierrez, Nehorai, & Preissl, 2005; J. G. Samuelsson, Peled, Mamashli, Ahveninen, & Hämäläinen, 2021). In the cerebellum, comprising about 80% of all neurons in the brain, equivalent sources were modeled as Purkinje cells, which possess large, planar dendritic trees oriented perpendicular to the local cerebellar surface (J. G. Samuelsson et al., 2020; Sereno et al., 2020). Since Purkinje cells constitute the sole output of the cerebellar cortex, they provide a physiologically appropriate basis for source modeling. The same orientation constraint—perpendicular to the local tissue surface—was applied to both cerebrum and cerebellum, enabling direct comparison of SNR across regions within a unified forward modeling framework.

A three-layer BEM model was generated with conductivities of 0.3, 0.006, and 0.3 S/m for the brain, skull and scalp, respectively. The cerebral surface was reconstructed using about 100,000 vertices per hemisphere, corresponding to an average vertex spacing of ∼ 2 mm.

### 2.4 ARCUS for cerebellar reconstruction and segmentation

ARCUS (John G Samuelsson et al., 2020) is an automated algorithm for cerebellar segmentation and surface reconstruction (Fig. 1). It employs subcortical segmentation surfaces extracted from FreeSurfer to isolate the cerebellum, followed by non-linear diffeomorphic registration of a high-resolution ex vivo cerebellar template to the subject’s MRI. This template-driven morphing enables approximate reconstruction of the cerebellar cortical surface even at typical clinical MRI resolutions (1 mm isotropic), which is insufficient for direct folium-level segmentation.

**Fig. 1.**
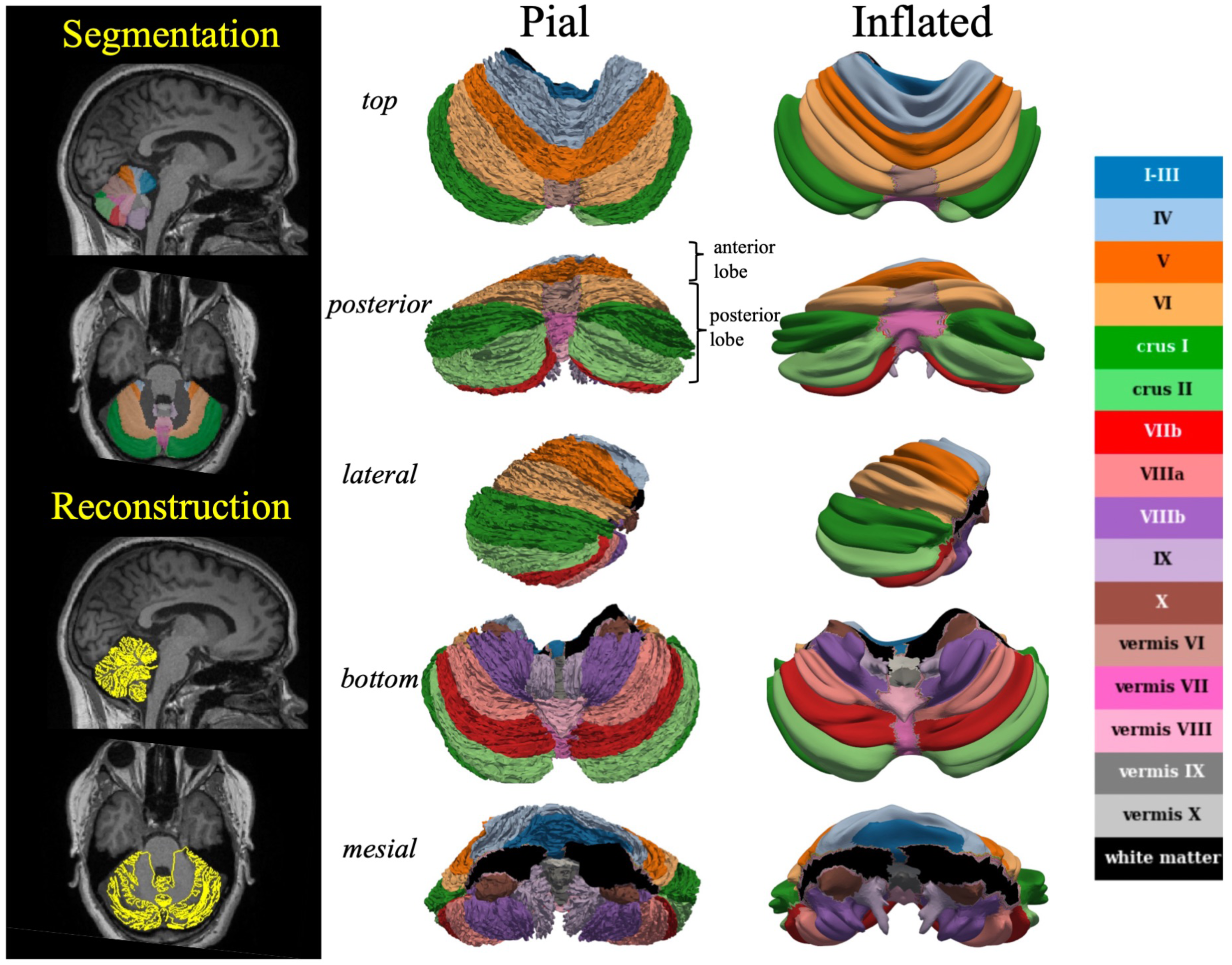
ARCUS-derived segmentation and reconstruction. Representative example of ARCUS-derived cerebellar reconstruction and lobular segmentation from clinical MRI. Colors denote cerebellar regions and are used consistently across all subsequent figures.

Briefly, the method applies an affine alignment of the ex vivo template to the subject’s cerebellar volume, followed by nonlinear registration using symmetric normalization with cross-correlation (SyNCC, ANTs implementation). The resulting deformation fields are applied to the template surface to generate a subject-specific cerebellar mesh, with vertex normal recalculated in native MRI space.

In this study, ARCUS was applied within an MNE-Python-compatible workflow using dense spatial sampling, yielding 99,856 vertices—approximately matching the vertex density of one cerebral hemisphere.

One of the current study’s objectives was to evaluate the feasibility of applying ARCUS to clinical MRI data of heterogeneous quality. To ensure anatomical plausibility at the spatial scale relevant for this study, ARCUS-derived cerebellar segmentations and source-space representations were visually inspected by an experienced neurophysiologist and neurosurgeon (T.M.). Visual inspection focused on the consistency of lobular boundaries and overall cerebellar morphology with individual MRI, rather than on folium-level surface accuracy.

### 2.5. Signal-to-noise ratio (SNR) for individual sources

Previous on-scalp MEG studies have primarily characterized sensitivity at the sensor or vertex level, or in terms of field-of-view and information capacity, focusing on the cerebral cortex (Hill et al., 2020; Iivanainen et al., 2017). While these approaches have provided important system-level insights, they have not quantified sensitivity at the level of anatomically defined brain regions, nor have they investigated the cerebellum in a population-based framework. In the present study, we extend system-level SNR modeling to anatomically defined regions of both the cerebrum and cerebellum, enabling region-wise comparison of signal detectability across modalities and sensor configurations in a large clinical cohort.

SNR was quantified following the formulation of Goldenholz et al. (Goldenholz et al., 2009), extended here to include cerebellar sources and simulated OPM configurations. For each source location, the SNR was defined in decibel (dB) units as:

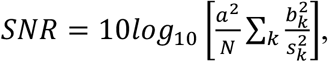

where *b_k_* denotes the signal at sensor *k* provided by the forward model for a unit-amplitude source, *a* is the assumed source amplitude, *N* is the number of sensors, and 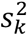 is the noise variance at sensor *k*. The unit source amplitude was fixed at 10 nAm based on the Okada constant (Murakami & Okada, 2015) and applied uniformly across all sources and modalities. This formulation yielded an SNR estimate for each vertex across both cerebral and cerebellar source spaces. Vertex-wise SNR reflects the combined effects of source-sensor distance, source orientation, and head geometry encoded in the forward model, together with the assumed noise structured described in Section 2.6.

For cross-modality comparisons, SNR difference was defined as the vertex-wise subtraction of SNR values between modalities (in dB), with positive values indicating higher SNR in the first modality.

Since OPM recordings were not available from patients in this cohort, OPM SNR was estimated by simulation using subject-specific forward models generated under multiple sensor-configuration conditions (Section 2.7).

#### 2.6.1. Noise estimation

Noise variance estimation followed the procedure described by Goldenholz et al (Goldenholz et al., 2009). Specifically, the noise variance 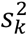 for each sensor was estimated from spontaneous clinical MEG and EEG recordings. Sixty seconds of resting-state data were extracted and band-pass filtered between 0.5 Hz and 100 Hz. Five patients were excluded from this analysis and all subsequent analyses because frequent interictal spikes during the resting recording prevented reliable estimation of baseline noise.

Following the Goldenholz framework, the noise variance at sensor *k* was modeled as:

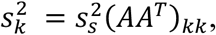

where *A* denotes the forward-solution matrix for unit-amplitude noise sources and 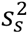 represents the variance of cortical source activity. This model assumes a spatially uniform distribution of independent noise sources, i.e., equal strength in all vertices of the source-space, oriented perpendicular to the cortical surface, with Gaussian-distributed amplitudes of zero mean and variance 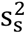. Instrumentation noise was not explicitly modeled, such that the estimated noise variance reflects biologically plausible background brain activity as mediated by the forward model.

For each modality, median normalized values of 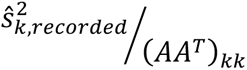 were computed separately for SQUID magnetometers, SQUID gradiometers and EEG electrodes. The weighted mean of these three medians, proportional to the number of sensors in each category, was taken as the overall estimate of 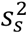. This procedure accounts for realistic brain-noise contributions while minimizing bias from individual bad channels. The resulting brain-noise estimate was applied consistently for the computation of SNR in SQUID-MEG, EEG, and simulated OPM configurations. Validation supporting the application of this common noise estimate to OPM simulations is described in Section 2.6.2.

#### 2.6.2 OPM noise estimation

To validate the use of OPM noise levels estimated from SQUID and EEG data in individual subjects, we compared OPM and SQUID data acquired from a single healthy participant. OPM recordings (FieldLine, Gen1) were obtained in a single-layer magnetically shielded room, whereas SQUID recordings were acquired on the same day from the same subject in a three-layer magnetic shield room at the Martinos Center. During OPM acquisition, biplanar field-nulling coils were used to compensate residual magnetic fields, reducing them to approximately 4 nT.

The SQUID system comprised a whole-head 306-channel array (102 magnetometers and 204 gradiometers). The OPM system consisted of a helmet-type array with 16 sensors primarily covering the left somatosensory region. While the subject was seated comfortably, electrical stimulation was applied to the right median nerve to evoke somatosensory responses. Only magnetometers channels were analyzed. Two of the sixteen OPM sensors were excluded due to poor signal quality.

Both datasets were band-pass filtered between 2 Hz and 100 Hz, and a 60 Hz notch filter was applied. For OPM data, homogeneous field correction (Tierney et al., 2021) was used to suppress residual homogeneous environmental noise within the shielded room. For SQUID-MEG data, signal-space projection (Uusitalo & Ilmoniemi, 1997) based on empty-room recordings was applied for the same purpose. Effectively the environmental magnetic interferences were removed in both modalities, minimizing the instrumentation noise.

After equalizing the number of trials (338), averaged evoked responses were computed (Supplementary Fig. 1A, B). Baseline activity from -200 ms to -5 ms before stimulus onset was used to estimate sensor-level noise variance for each sensor. To enable a fair comparison between SQUID and OPM sensors with different forward sensitivities, sensor-level noise variance was normalized by the corresponding forward-model sensitivity, thereby expressing noise in units relative to source-space detectability. Statistical comparison between modalities was performed on these normalized noise measures, which reflect the relative contribution of individual sensors to source detectability, using Welch’s t-test. In addition, nonparametric bootstrap resampling (1000 iterations) was used to estimate the 95% confidence interval of the mean difference. For each bootstrap iteration, values were sampled with replacement from both datasets and the difference in sample means was computed. The 2.5^th^ and 97.5^th^ percentiles of the resulting bootstrap distribution define the confidence interval. Effect size was quantified using Cohen’s d to assess the magnitude of the difference between modalities.

No significant difference in noise levels was observed between OPM and SQUID recordings (Supplementary Fig. 1C; t = -0.35, p = 0.72). The 95% confidence interval for the mean difference was [-3.02e-22, 1.65e-22], encompassing zero, and the effect size was negligible (Cohen’s d = -0.06). These findings indicate that the estimated brain-noise levels were comparable between OPM and SQUID magnetometer sensor classes, supporting the validity of treating OPM sensors as equivalent to SQUID magnetometers within the same brain-noise estimation framework. This, in turn, justifies applying the brain-noise estimates derived from the SQUID-EEG recordings to the OPM simulations used in this study.

### 2.7. OPM configurations

Because OPM recordings were not available from individual patients with epilepsy, multiple simulated OPM configurations were generated to estimate SNR for both the cerebrum and cerebellum, and to enable comparison with clinical MEG (Fig. 2A; SQUID_clinical_) and clinical EEG (Fig. 2B; EEG_clinical_). For consistency, all simulations used FieldLine Gen1 sensors, identical to those used in the real OPM recordings described in Section 2.6.2.

**Fig. 2.**
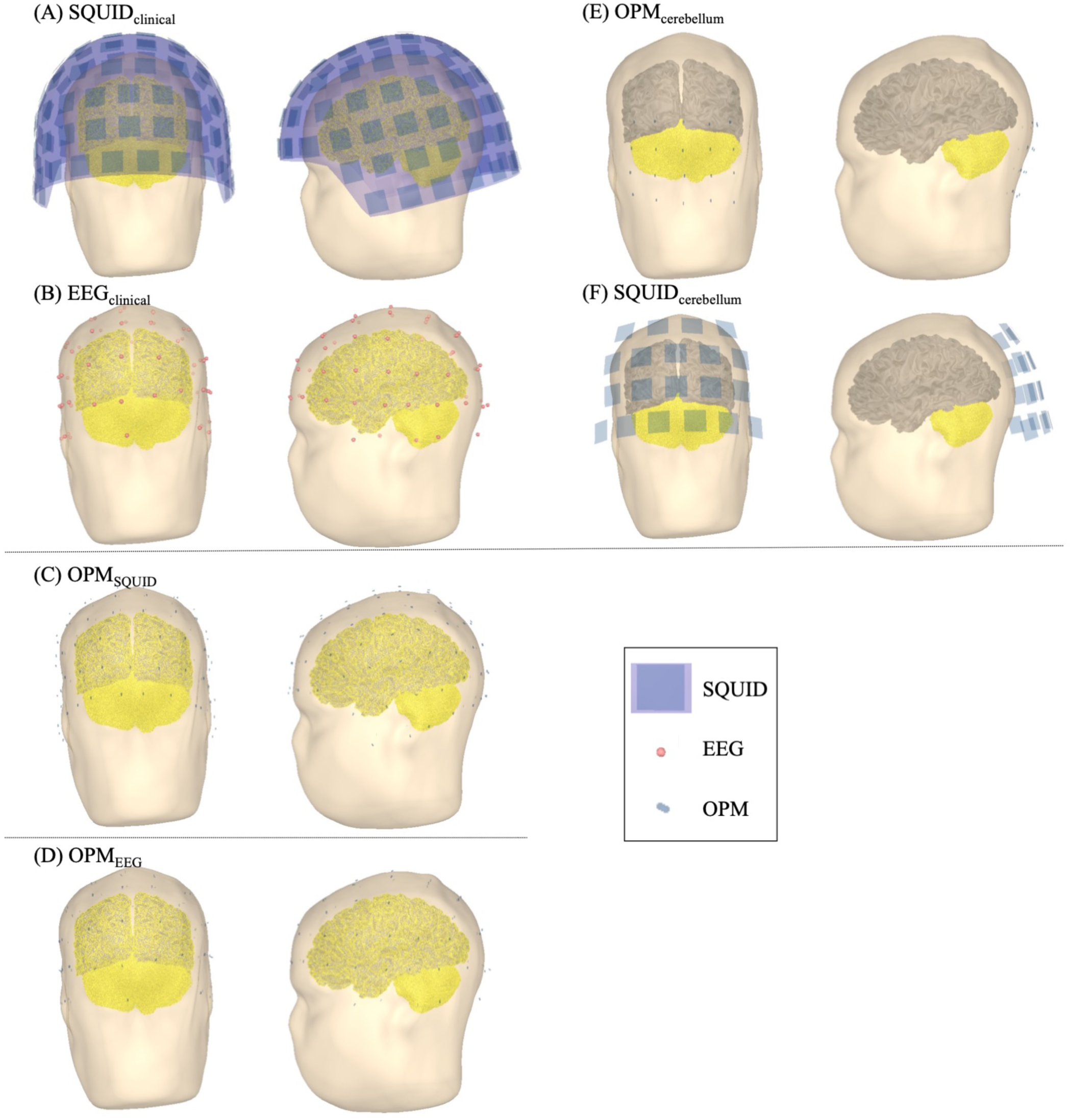
Sensor configurations. (A) SQUID_clinical_: Clinical whole-head SQUID system consisting of 306 channels (102 magnetometers and 204 gradiometers). (B) EEG_clinical_: Clinical EEG system using 70 scalp electrodes. (C) OPM_SQUID_: Simulated OPM layout with sensors positioned at scalp locations corresponding to the clinical SQUID array projected onto the head surface with a 5 mm offset. (D) OPM_EEG_: Simulated OPM layout with sensors positioned at scalp locations corresponding to the clinical EEG electrode positions. (E) OPM_cerebellum_: Cerebellar-optimized OPM layout consisting of a 5 × 4 grid (10 by 8 cm) placed over the occipital region, with the superior edge aligned to the inion. (F) SQUID_cerebellum_: SQUID layout selected for comparison with OPM_cerebellum_, consisting of 20 fixed SQUID sensors covering the occipital region.

Three OM configurations were generated:

i. OPM_SQUID_ (Fig. 2C): To enable a comparison with the clinical SQUID system, 102 virtual OPM sensors were positioned at locations corresponding to SQUID magnetometer positions projected onto the scalp surface, with a fixed 5 mm offset. SNR difference was computed as the subtraction between source-space SNRs observed from OPM_SQID_ and SQUID_clinical_.
ii. OPM_EEG_ (Fig. 2D): To compare OPM and EEG under equivalent spatial coverage, 70 OPM sensors were placed at the same scalp locations as standard EEG electrodes based on Polhemus digitization. This configuration enabled direct comparison between OPM and EEG channels using on-scalp sensor placement, with source-space SNR difference between OPM_EEG_ and EEG_clinical_ (defined as OPM_EEG_ minus EEG_clinical_ SNR).
iii. OPM_cerebellum_ and SQUID_cerebellum_ (Fig. 2E, F): Whereas the first two configurations reflect conventional clinical layouts optimized for cerebral coverage, this configuration was designed specifically to enhance posterior fossa and cerebellar sensitivity. Twenty OPM sensors were arranged in a 5 × 4 grid (10 cm × 8 cm) over the occipital region. The upper edge of the grid was aligned with the inion, as identified on each subject’s MRI, to anchor the array anatomically relative to the cerebellum. For fair comparison, an equivalent SQUID_cerebellum_ configuration was defined by selecting twenty fixed SQUID sensors covering the occipital region. Because SNR was normalized by sensor count, the same number of sensors was used for both systems. The SNR difference was computed between OPM_cerebellum_ and SQUID_cerebellum_. Analyses for this configuration were restricted to cerebellar regions.

### 2.8. Source orientation (folded angle)

To investigate whether regional variability in source orientation contributes to differences in signal detectability between EEG and MEG comparison, we analyzed the relationship between local source orientation and SNR difference in the OPM_EEG_-EEG_Clinical_ comparison scenario. For each subject and vertex, source orientation was quantified as the angle between the surface normal vector and a vector extending from the center of the inner skull surface toward that vertex. Angles were folded into the 0–90° range, where 90° corresponds to a tangential orientation relative to the skull surface (generally favorable for MEG detection), and 0° corresponds to radial orientation.

### 2.9. Sensor distance ratio

To evaluate how anatomical and geometric factors contribute to SNR differences between OPM_cerebellum_ and SQUID_cerebellum_, we quantified the relative proximity of sensors to cerebellar sources. For each cerebellar vertex, the shortest Euclidean distance to both OPM and SQUID sensors was computed. The sensor distance ratio was defined as the SQUID distance divided by the OPM distance, such that larger values indicate proximity advantage for OPM sensors. Since OPM sensors are positioned closer to the scalp compared to SQUID sensors, the distance ratio was consistently greater than one across all vertices.

### 2.10. Statistical analysis

#### 2.10.1. Region of interest

For each metric (SNR, SNR difference, folded angle, and sensor distance ratio), values were averaged within regions of interest (ROIs), combining left and right hemispheres. The sensor distance ratio was only computed for cerebellar ROIs. Regional SNR difference was tested against zero using the Wilcoxon signed-rank test, with false discovery rate (FDR) correction applied separately for cerebrum and cerebellum. Statistical significance was assessed using a two-sided alpha of 0.05 after correction.

For the cerebrum, ROIs were defined to provide a reference framework for cerebellar comparisons rather than a detailed cortical analysis. Accordingly, broad cortical regions were used: frontal, lateral temporal, medial temporal, parietal, occipital, cingulate, and insular regions. These labels were derived from FreeSurfer Desikan-Killiany cortical parcellation. The medial temporal ROI was constructed by combining entorhinal, parahippocampal, fusiform and temporal pole regions.

For the cerebellum, a full set of ARCUS-derived ROIs was analyzed to enable detailed regional evaluation. These included lobules I-III, IV, V, VI, Crus I, Crus II, VIIb, VIIIa, VIIIb, IX, X, as well as vermis VI, vermis VII, vermis VIII, vermis IX, and vermis X.

#### 2.10.2 Distance

To investigate anatomical and geometric contributions to SNR differences between OPM_cerebellum_ and SQUID_cerebellum_ comparison scenario, three variables were examined across cerebellar regions: (A) SNR difference, defined in Section 2.7.iii as the regional cerebellar SNR of OPM_cerebellum_ minus that of SQUID; (B) Sensor distance ratio, defined in Section 2.9.; and (C) Head circumference, used as a proxy for lateral head size.

Pairwise Pearson correlations and linear regression analyses were first performed to characterize direct relationships among these values. Head circumference was negatively correlated with the sensor distance ratio (Fig. 5D), indicating that in individuals with larger heads, the relative proximity of OPM and SQUID sensors become more similar. A positive correlation was observed between the distance ratio and the SNR difference (Fig. 5E), consistent with greater SNR advantage for OPM when sensors are relatively closer to the cerebellum.

To determine whether the effect of the head size on SNR was mediated by the sensor distance ratio, mediation analysis (Baron & Kenny, 1986) was performed for each cerebellar ROI. Specifically, we estimated:

a. The a-path, representing the effect of head circumference on the distance ratio.
b. The b-path, representing the effect of the distance ratio on SNR difference after controlling for head circumference; and
c. The indirect effect (a × b), representing the mediated influence of head size through relative sensor distance.

Statistical significance of indirect effects was evaluated using Sobel’s test, with FDR correction applied across cerebellar ROIs.

#### 2.10.3 Source orientation

To evaluate whether source orientation contributed the regional SNR patterns in the OPM-EEG comparison (OPM_EEG_ vs EEG_Clinical_) where sensor geometry was identical, linear regression was applied using all pooled data across subjects and regions. Folded angle was regressed against SNR difference, and statistical significance of the regression slope was assessed using SciPy’s linregress function.

## 3. Results

### 3.1 ARCUS feasibility on clinical MRI data

ARCUS processing was performed in 62 patients. Visual inspection confirmed anatomically plausible cerebellar segmentation in 59 of 62 patients (success rate of 95.1%). Representative examples of successful segmentations and all failure cases are shown in Supplementary Figs. 2, and 3, with a summary provided in Table 1.

**Table 1.**
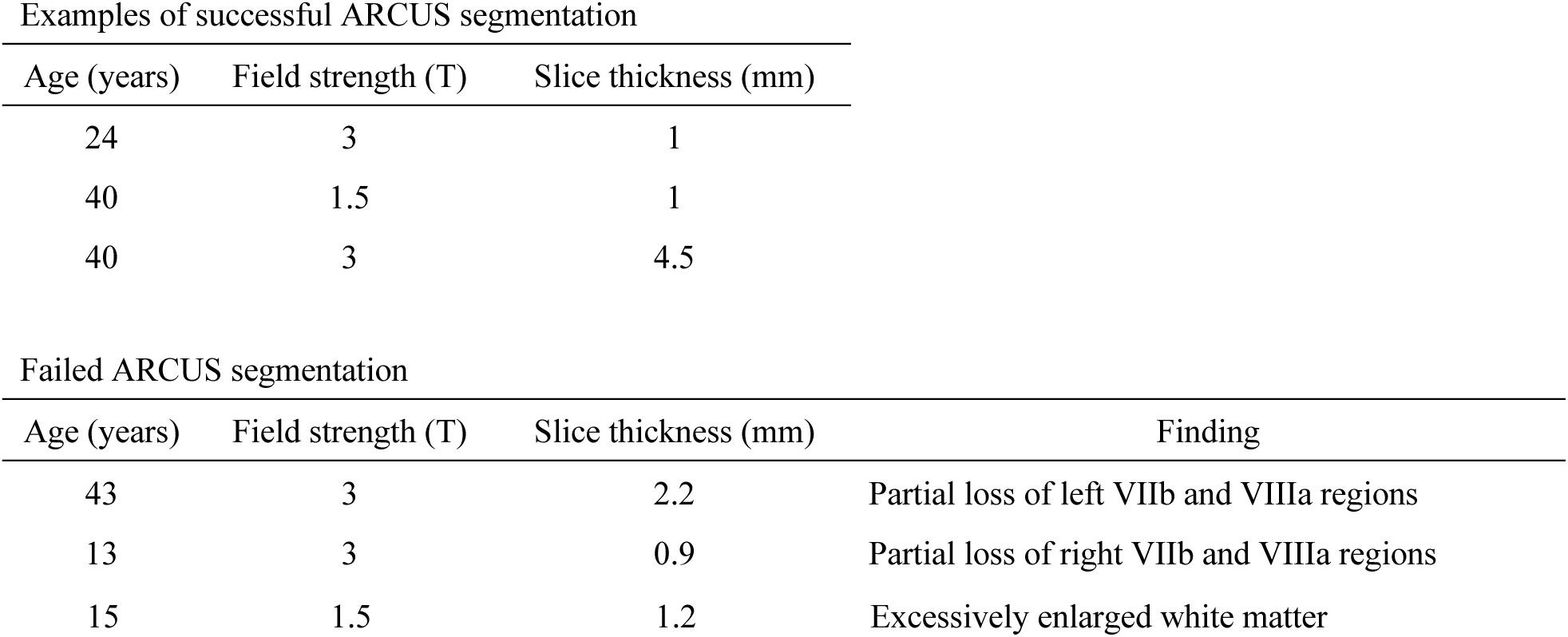
Examples of successful ARCUS segmentation (Supple Fig. 2) and failed cases (Supple Fig. 3)

Successful segmentations were obtained even in several cases acquired at 1.5 T or with relatively thick slice thicknesses (up to 4.5 mm). In contrast, the failed cases were characterized by poor tissue contrast, particularly reduced distinction between cerebellar cortex and white matter. In one failed 1.5 T case, this blurring likely contributed to loss of cerebellar surface definition.

Overall, the high success rate across heterogeneous clinical MRI datasets supports the feasibility of applying ARCUS to routine clinical imaging for cerebellar source modeling. For subsequent analyses, five patients were excluded due to unreliable noise estimation (Section 2.6.1), resulting in a final sample of 54 patients.

### 3.2 SNR across modalities

#### 3.2.1. SQUID_clinical_ vs. EEG_clinical_

For the cerebrum, the median SNR in SQUID_clinical_ ranged from -32 dB to -23 dB (Fig. 3A). The highest SNR was observed in the parietal lobe (-23 dB, dashed line), consistent with close proximity of superficial cortical sources positioned close to the sensors. Regional SNR patterns followed cortical folding geometry, with higher SNR along sulcal walls—where MEG is most sensitive to tangential sources—and lower SNR in deeper structures, including the interhemispheric and sylvian fissure. In contrast, EEG_clinical_ showed less regional variability across the cerebellum (Fig. 3B), resulting in a more uniform SNR distribution. The SNR difference revealed that only the parietal lobe favored SQUID_clinical_, whereas most other regions—including the cingulate and insular–favored EEG (Fig. 3E), consistent with EEG’s greater sensitivity to deeper sources.

**Fig. 3.**
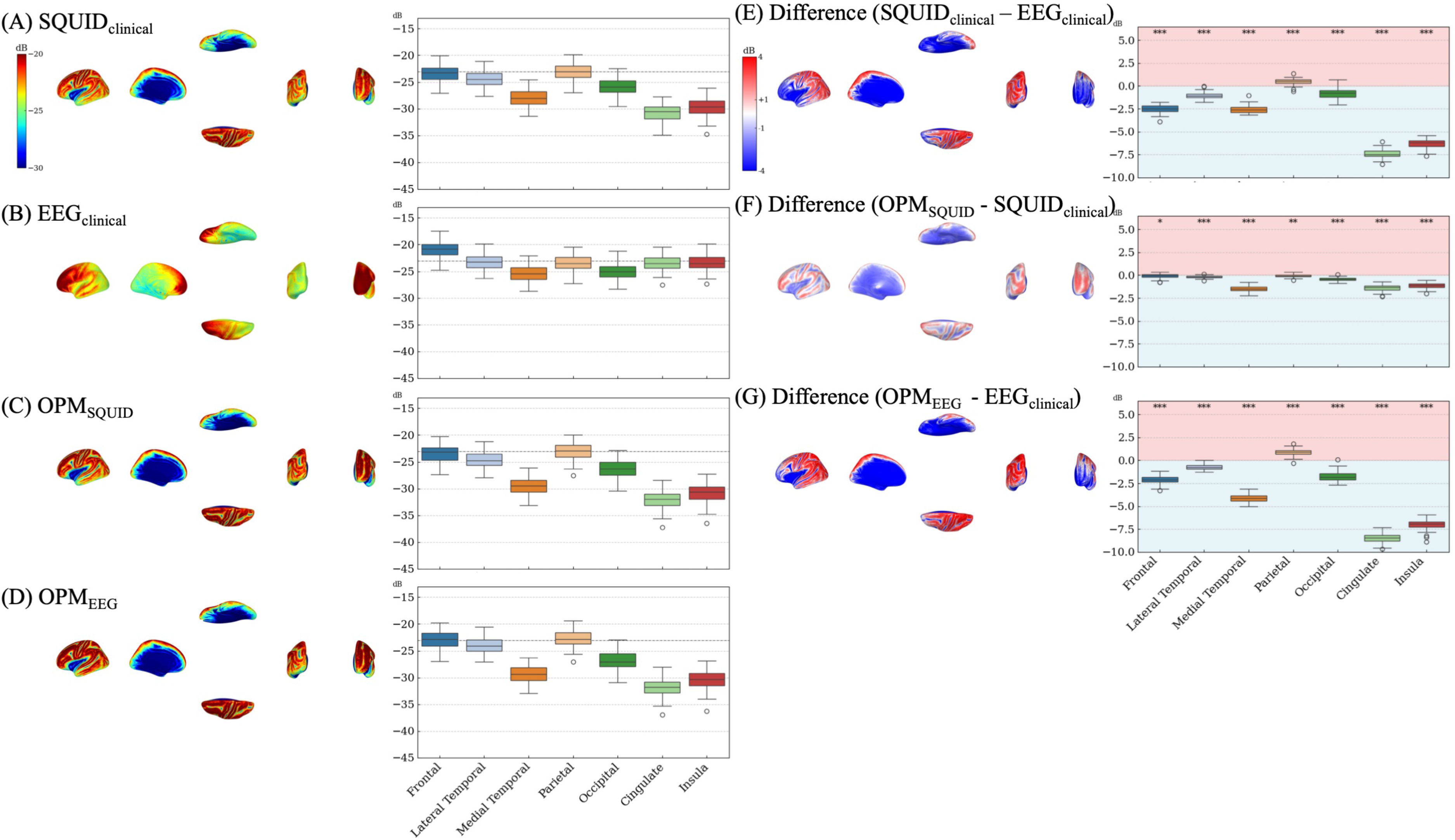
Cerebral SNR and SNR difference across modalities. Cerebral SNR maps for (A) SQUID_clinical,_ (B) EEG_clinical_, (C) OPM_SQUID_, and (D) OPM_EEG_. SNR difference maps comparing (E) SQUID_clinical_ vs. EEG_clinical_, (F) OPM_SQUID_ vs. SQUID_clinical_, and (G) OPM_EEG_ vs. EEG_clinical_. Asterisks indicate statistically significant SNR differences after false discovery rate (FDR) correction (* < 0.05, ** < 0.01, ***< 0.001). The boxplots show the SNR for each region, in the following order: frontal, lateral temporal, medial temporal, parietal, occipital, cingulate and insula.

For the cerebellum, median SNR in both SQUID_clinical_ (Fig. 4A), and EEG_clinical_ (Fig. 4B) remained below the -23 dB level corresponding to superficial cerebral cortex, indicating limited detectability in the clinical setup. The SNR difference showed that Crus I, Crus II, lobules VIIb, VIIIa, VIIIb, IX and X exhibited relatively higher SNR in SQUID_clinical_ compared with EEG_clinical_ (Fig. 4E).

**Fig. 4.**
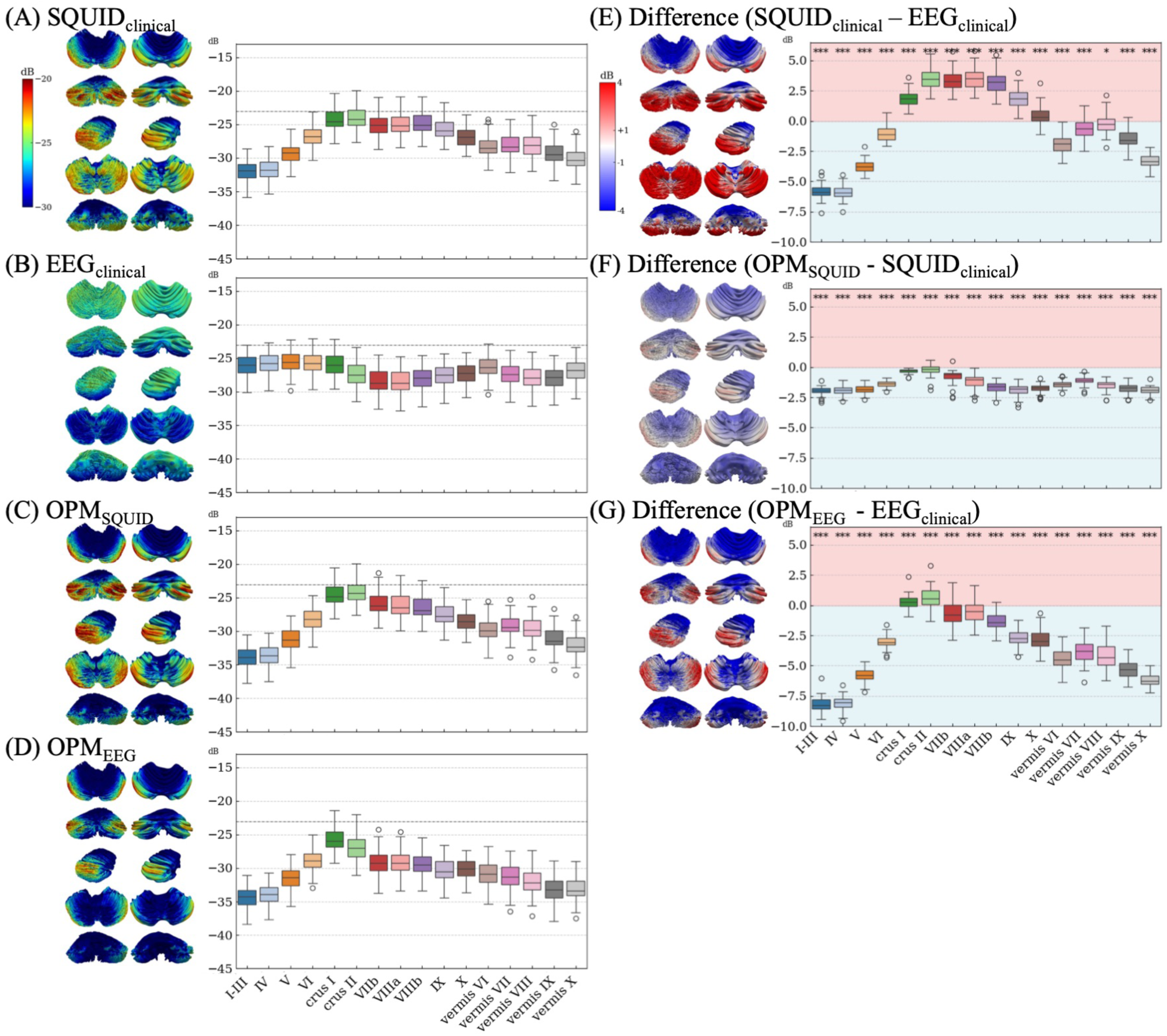
Cerebellar SNR and SNR difference across modalities. Cerebellar SNR maps for (A) SQUID_clinical,_ (B) EEG_clinical_, (C) OPM_SQUID_, and (D) OPM_EEG_. SNR difference maps comparing (E) SQUID_clinical_ vs. EEG_clinical_, (F) OPM_SQUID_ vs. SQUID_clinical_, and (G) OPM_EEG_ vs. EEG_clinical_. Asterisks indicate statistically significant SNR differences after FDR correction (* < 0.05, ** < 0.01, ***< 0.001). The boxplots show the SNR for each region, in the following order: lobules I-III, IV, V, VI, Crus I, Crus II, VIIb, VIIIa, VIIIb, IX, X, vermis VI, VII, VIII, IX, and X.

#### 3.2.2. OPM_SQUID_ vs. SQUID_clinical_

In OPM_SQUID_ configuration, OPM sensors were virtually positioned at the scalp locations corresponding to SQUID magnetometer positions projected onto the scalp surface, allowing comparison under the same acquisition modality while reducing sensor-source distance. For both the cerebrum and the cerebellum, the SNR maps from OPM_SQUID_ (Fig. 3C and Fig. 4C) closely resembled those from SQUID_clinical_ (Fig. 3A and Fig. 4A). The SNR difference showed only minimal differences, with no regions demonstrating higher for OPM_SQUID_ in either structure (Fig. 3F and Fig. 4F). These results indicate that merely reducing the sensor distance, while preserving geometry and coverage, does not yield meaningful SNR improvement for deeper sources.

#### 3.2.3. OPM_EEG_ vs. EEG_clinical_

In the OPM_EEG_ configuration, OPM sensors were placed at the same scalp locations as clinical EEG electrodes, enabling direct comparison under equivalent spatial coverage with on-scalp sensor placement. The results (Fig. 3D, 3G, 4D, 4G) mirrored those observed in the SQUID_clinical_–EEG_clinical_ (Section 3.2.1): only superficial regions, such as the parietal lobe and Crus I/II, showed higher SNR in OPM_EEG_.

#### 3.2.4. OPM_cerebellum_ vs. SQUID_cerebellum_

Figure 5A–C shows the comparison between OPM_cerebellum_ and SQUID_cerebellum_ using layouts specifically optimized for posterior fossa coverage. Since this configuration targets the cerebellum, analyses were restricted to cerebellar regions. Compared with the clinical configuration (Fig. 4A), the OPM_cerebellum_ layout yielded severalposterior regions—Crus I, Crus II, lobules VIIb, VIIIa, VIIIb—with SNR values exceeding the -23 dB superficial cerebral reference level (Fig. 5A). The SNR difference (Fig. 5C) further showed that these regions except for VIIIb, as well as vermis VII and vermis VIII exhibited higher SNR, favoring OPM.

**Fig. 5.**
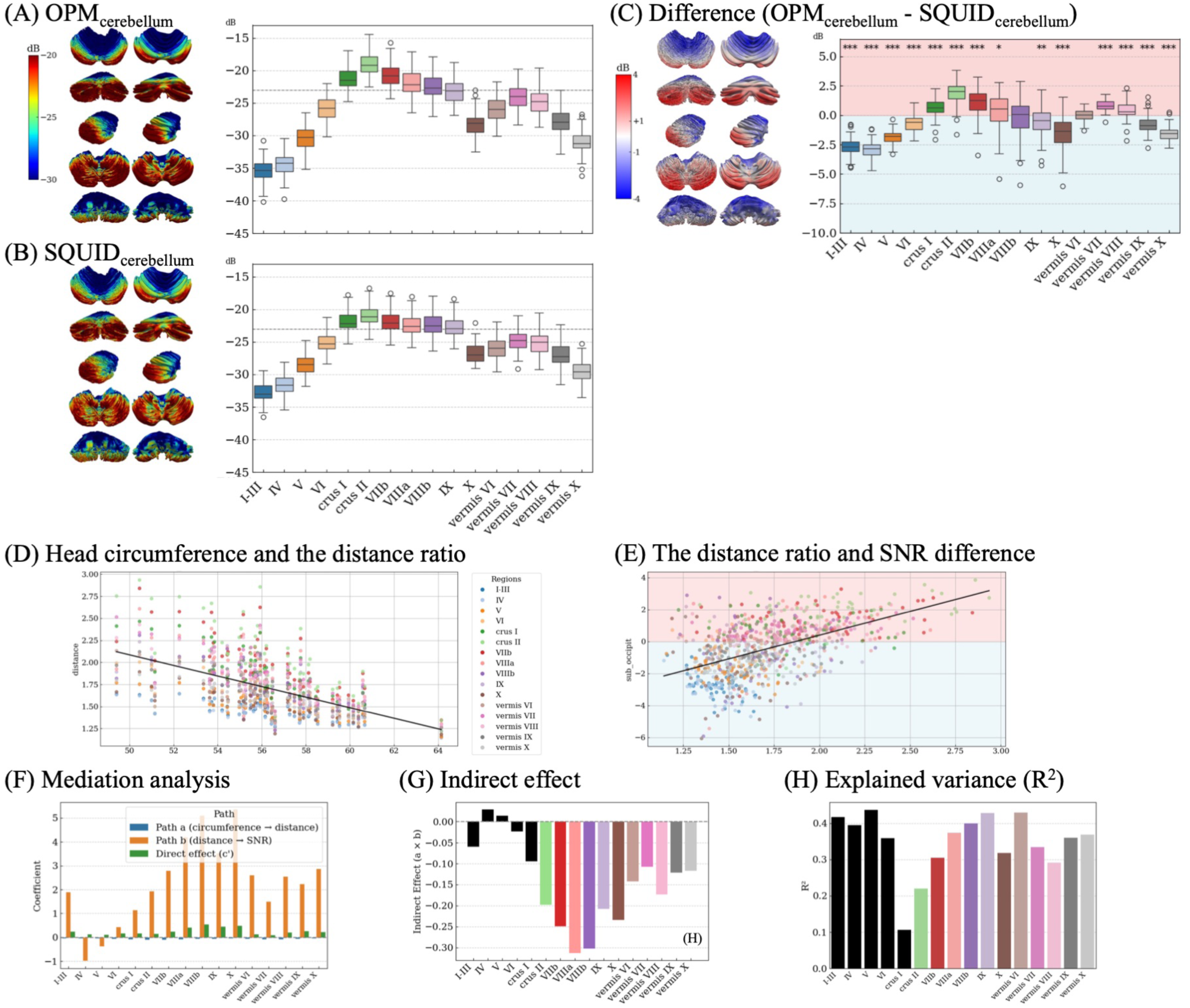
Cerebellar-optimized comparison between OPM_cerebellum and_ SQUID_cerebellum_. (A) SNR for OPM_cerebellum_ and (B) SNR for SQUID_cerebellum_. (C) SNR difference between OPM_cerebellum_ and SQUID_cerebellum_. (D) Relationship between head circumference and sensor distance ratio, showing a negative relationship. (E) Relationship between sensor distance ratio and SNR difference, showing a positive relationship. In panels, (D) and (E), each point represents a cerebellar region, and the solid line indicates the linear regression fit. (F) Mediation analysis showing the a-path (head circumference -> distance ratio), b-path (distance ratio -> SNR difference), and direct effect. (G) Indirect path (a × b) across regions. (H) Explained variance (R^2^) of the mediation model across cerebellar regions. Black bins indicate regions that did not reach statistical significance after FDR correction. Asterisks indicate statistical significance after FDR correction (* < 0.05, ** < 0.01, ***< 0.001). The boxplots show the SNR for each region, in the following order: lobules I-III, IV, V, VI, Crus I, Crus II, VIIb, VIIIa, VIIIb, IX, X, vermis VI, VII, VIII, IX, and X.

### 3.3 Distance effect on SNR for OPM_cerbellum_ and SQUID_cerebellum_

Across cerebellar regions, head circumference showed a negative correlation with sensor distance ratio (Fig. 5D; r = -0.56, p < 0.001), indicating that larger head size reduced the relative proximity advantage of OPM sensors. The sensor distance ratio was positively correlated with SNR difference (Fig. 5E; r = 0.53, p < 0.001), consistent with greater SNR benefit when OPM sensors were relatively closer to the cerebellum.

Mediation analysis (Fig. 5F) revealed significant indirect effects (a × b) in most cerebellar regions after FDR correction, with the exception of the anterior lobes (I-III, IV, V) and Crus I, (Fig. 5G). These findings indicate that the head size influences cerebellar SNR difference primarily through its effect on relative sensor distance, particularly in posterior regions where OPM advantage was largest.

Direct effects of head circumference on SNR difference (c’ path), after controlling for distance, were generally small or non-significant, consistent with distance acting as the primary mediator of head size effects on signal detectability.

Some regions such as I-III, IV, V, VI, and Crus I did not show statistically significant indirect effects after FDR correction, despite relatively high R2 values in the regression models (Fig. 5H). Notably, the b-path was negative in a few regions (e.g., IV and V), which deviates from the expected direction. While these may reflect local anatomical or modeling differences, their non-significance after correction suggests that we cannot draw firm conclusions from these specific deviations.

Overall, these results support a mediated relationship where individuals with smaller heads (greater distance asymmetry between OPM and SQUID) tend to exhibit stronger SNR benefit from OPMs in posterior cerebellar regions.

### 3.4 Orientation effect on SNR difference for OPM_EEG_-EEG_clinical_

Folded angle maps for the cerebrum and cerebellum are shown in Fig. 6A and 6B, together with scatter plots illustrating the relationship between folded angle and SNR difference. Substantial intersubject variability in folded angle is observed, particularly in the medial temporal lobe in the cerebrum (Fig. 6A), as well as across most cerebellar regions (Fig. 6B).

**Fig. 6.**
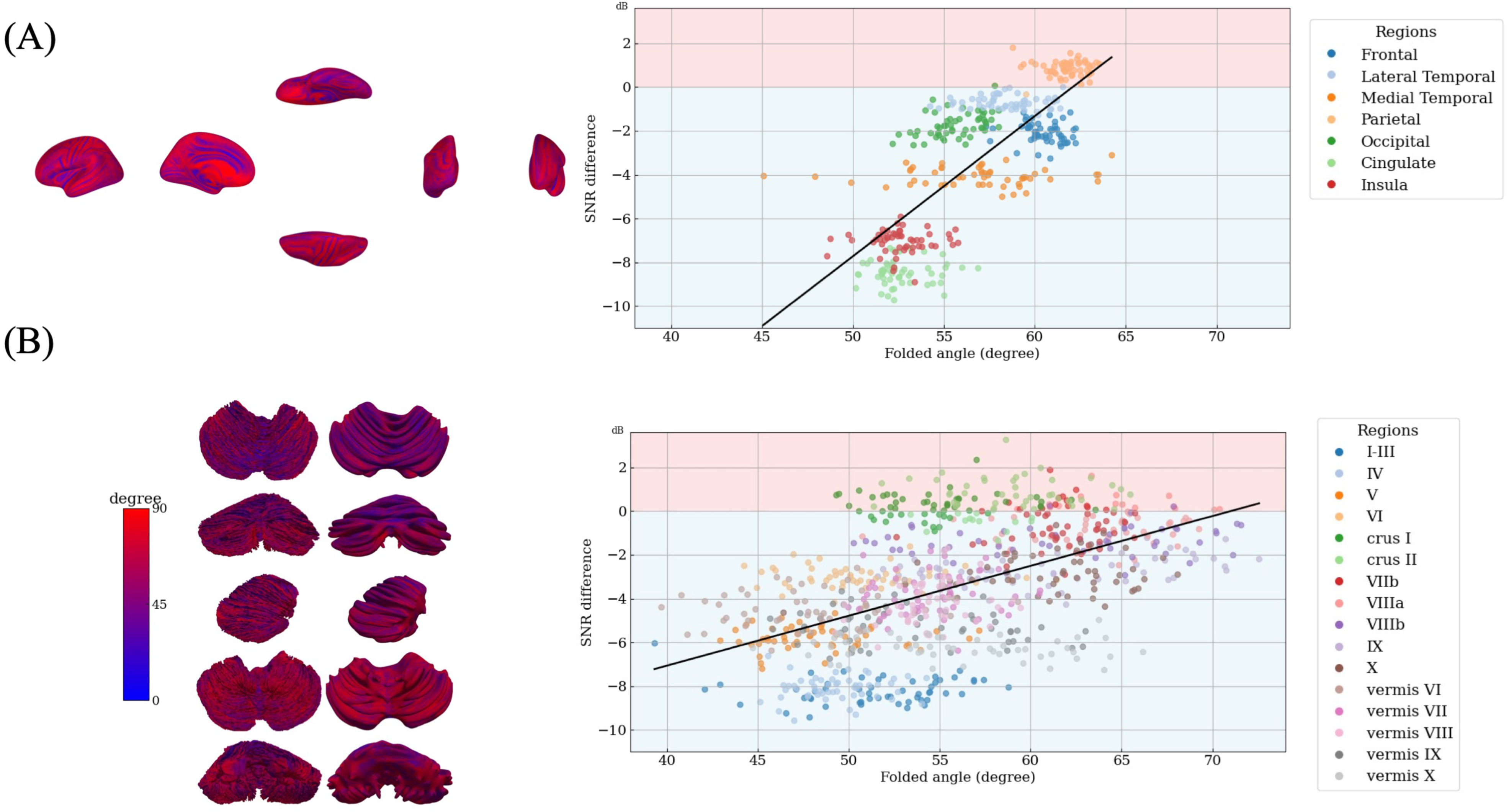
Orientation effects on SNR difference in the OPM_EEG_ configuration. Folded angle maps for the cerebrum (A) and cerebellum (B) in the OPM_EEG_ layout. Right panels show scatter plots illustrating the relationship between folded angle and SNR difference across regions. Solid lines indicate linear regression fits.

Linear regression analysis demonstrated a stronger relationship between folded angle and SNR difference in the cerebrum (slope = 0.64, p < 0.001) than in the cerebellum (slope = 0.23, p < 0.001). These results indicate that source orientation contributes more strongly to detectability differences in the cerebrum, whereas in the cerebellum, depth and geometric dispersion dominate OPM sensors’ sensitivity, when OPM sensors are placed at clinical EEG electrode positions.

## 4. Discussion

In this study, we combined individualized cerebellar modeling using ARCUS with modality-specific forward modeling to evaluate signal detectability in the cerebellum across clinical SQUID-MEG, EEG and simulated OPM configurations in a cohort of patients with epilepsy. By quantifying SNR and SNR difference across modalities and sensor layouts, we provide a population-level assessment of how cerebellar anatomy, source geometry, and sensor configuration jointly shape the sensitivity of various noninvasive electrophysiological measurements across different modalities. Our results demonstrate that cerebellar detectability is strongly constrained in routine clinical settings by the neuroanatomy, but can be selectively enhanced through cerebellar-optimized OPM configurations, particularly in posterior lobes.

### 4.1 Cerebellar functional organization and relevance to epilepsy

The cerebellum exhibits a well-established functional organization along the anterior-posterior axis. In the healthy human brain, anterior lobes are predominantly associated with sensorimotor processing (Grodd, Hulsmann, Lotze, Wildgruber, & Erb, 2001; Schmahmann & Sherman, 1998), whereas posterior lobules are more strongly linked to high-order cognitive and affective functions (Buckner, Krienen, Castellanos, Diaz, & Yeo, 2011; King, Hernandez-Castillo, Poldrack, Ivry, & Diedrichsen, 2019; Stoodley & Schmahmann, 2009). This large-scale functional differentiation provides a useful anatomical framework for interpreting region-specific cerebellar signals in electrophysiological studies, independent of disease context.

In epilepsy, most evidence for cerebellar involvement in human brain function has come from structural (Allen et al., 2019; Lawson, Vogrin, Bleasel, Cook, & Bye, 2000; Sandok, O’Brien, Jack, & So, 2000), metabolic (Bohnen, O’Brien, Mullan, & So, 1998; Laich et al., 1997; Seto et al., 1997; Shin, Hong, Tae, & Kim, 2002; Sone et al., 2022), and postmortem investigations (Crooks, Mitchell, & Thom, 2000). Despite this converging evidence, direct non-invasive electrophysiological characterization of cerebellar activity has remained limited (Okada et al., 2020), possibly due to technical constrains on signal detectability. This gap motivates the present focus on cerebellar SNR characteristics across modalities and sensor configurations.

### 4.2 ARCUS as a framework for cerebellar electrophysiology

There are many challenges in cerebellar electrophysiology (John G Samuelsson et al., 2020; J. G. Samuelsson et al., 2020). The primary challenge is that cerebellar foliation occurs at spatial scales comparable to, or smaller than, typical clinical MRI resolution, resulting in pronounced orientation dispersion and signal cancellation at the spatial distance relevant for MEG and EEG measurements. Under these conditions, a folium-resolved surface representation is not reconstructable from the resolution attainable with in vivo MRI, which is prerequisite for M/EEG source modeling. ARCUS was therefore developed to provide a geometrically informed reconstruction of the cerebellar cortex suitable for electrophysiological source modeling.

A key feature of ARCUS is the use of a high-resolution ex vivo cerebellar template to preserve the native foliation geometry that is otherwise unresolved in standard MRI. By diffeomorphically registering this template to individual anatomy, ARCUS provides a representation of the densely folded cerebellar cortex that preserves folding geometry at a scale relevant for electromagnetic forward modeling. Importantly, the ex vivo template is not used to impose subject-specific foliar detail beyond what is supported by the individual lobular segmentation data, but rather to inform a source-space representation that respects the intrinsic folding geometry of the cerebellar cortex. In this framework, vertices in the ARCUS-derived source-space should not be interpreted as representing individual folia or local cortical patches, but instead as discretization points of a distributed cortical geometry whose fine-scale folding is preserved in aggregate. This representation is well aligned with the biophysical reality that cerebellar electromagnetic signals arise from spatially distributed current sources whose net contribution reflects both geometry and orientation dispersion.

This approach fundamentally distinguishes ARCUS from atlas-based cerebellar parcellation methods such as SUIT or FreeSurfer-based cerebellar parcellations (Buckner et al., 2011; Diedrichsen, 2006; Diedrichsen, Balsters, Flavell, Cussans, & Ramnani, 2009; Diedrichsen et al., 2011; Faber et al., 2022; Han, Carass, He, & Prince, 2020; Lyu, Wu, Huynh, Ahmad, & Yap, 2024). While these methods provide anatomically defined, and more recently functionally defined (Nettekoven et al., 2024), lobule-level parcellations that are highly valuable for morphometric and volumetric analyses, they do not provide a cerebellar cortical reconstruction that adequately captures the cortical folding which can be used to model neuronal currents in Purkinje dendrites relevant for electromagnetic source modeling. In contrast, ARCUS provides an explicit cortical geometry derived from a high-resolution cerebellar template, enabling a source discretization that is directly informed by cerebellar folding and source organization, and is therefore naturally compatible with forward models used in MEG and EEG.

The theoretical feasibility of modeling cerebellar activity with M/EEG was first established in our earlier work (J. G. Samuelsson et al., 2020), which demonstrated that cerebellar activity can generate measurable electromagnetic signals despite substantial foliation-induced cancellation, using a high-resolution ex vivo cerebellar template. Building on this theoretical foundation, subsequent work showed that such high-resolution cerebellar surfaces can be adapted for electrophysiological source modeling in controlled research settings (Alho et al., 2023).

The present study extends these prior efforts by demonstrating that the ARCUS framework enables cerebellar source modeling robustly and at scale in heterogeneous real-world clinical MRI datasets. By systematically applying ARCUS to routine clinical MRI data acquired under variable imaging conditions, we show that anatomically valid cerebellar segmentations and corresponding source spaces can be obtained in the vast majority of patients (95%), despite variability in field strength and slice thickness. This robustness is essential for translating cerebellar electrophysiology from proof-of-principle studies to large-scale clinical and population-based investigations. Although we did not quantitatively assess folium-level reconstruction accuracy, visual inspection against individual MRI confirmed anatomically plausible lobular segmentation in the vast majority of cases, supporting the suitability of the derived source spaces for forward modeling relevant for the spatial scale considered here.

### 4.3 Cerebellar SNR across modalities and sensor configurations

Our SNR analyses suggest that, in routine clinical SQUID-MEG and EEG recordings, detection of cerebellar activity is challenging (Fig. 4A). When OPM sensors were simulated at scalp locations corresponding to the SQUID array—thereby reducing sensor-source distance while preserving geometry and coverage—no meaningful improvement in cerebellar SNR was observed (Fig. 4 F). In contrast, cerebellar-optimized OPM configurations yielded substantial SNR improvements in posterior cerebellar regions (Fig. 5C). Together, these results demonstrate that targeted coverage and geometry, rather than sensor proximity alone, are critical for improving detectability of deep and highly folded structures such as the cerebellum. Importantly, this pattern was not unique to the cerebellum. Several deep cerebral regions—including the medial temporal lobe, cingulate cortex, and _insulaalso_ failed to show meaningful SNR improvement when OPM sensors were placed closer to the scalp using SQUID-matched geometry (Fig. 3F). This convergence across cerebellar and cerebral regions provides and important insight: for deep sources, scalp-source distance is a dominant limiting factor, and reducing sensor distance by bringing sensors closer to the scalp alone does not significantly improve SNR (Bezsudnova, Koponen, Barontini, Jensen, & Kowalczyk, 2022; Boto et al., 2016; Iivanainen et al., 2017).

One plausible explanation is that moving sensors closer increases both signal and background brain-noise contributions in a comparable manner for deep sources, producing minimal net SNR gain (Brookes et al., 2022). In the present framework, SNR is determined by the forward model together with the assumed noise structure; therefore, differences in SNR across sensor layouts primarily reflect differences in the forward solution rather disproportionate enhancement of signal over noise. Under these conditions, simply shifting a fixed geometry closer to the scalp does not overcome the fundamental depth-related limitations.

By contrast, the cerebellar-optimized OPM configurations explicitly targets the region of interest by altering coverage, while modeling the effect of geometry via forward modeling, thereby improving alignment between sensor sensitivity profiles and cerebellar source distributions. The resulting SNR gains—most evident in posterior lobules—highlight that the principal advantage of OPMs lies not in proximity alone, but in the ability to design sensor layouts optimized for specific deep structures, which is not possible in fixed-helmet SQUID systems.

### 4.4 Orientation effects in the OPM-EEG comparison

Given the recognition that depth imposes a dominant constraint on detectability, we next examined the contribution of source orientation to SNR differences when OPM and EEG were compared under identical sensor coverage (Fig. 2B and 2D). In this configuration, folded angle emerged as a statistically significant determinant of SNR difference, with different magnitudes in the cerebrum and cerebellum.

The relationship between folded angle and SNR difference was stronger in the cerebrum (Fig. 6A) than in the cerebellum (Fig. 6B). Notably, the medial temporal lobe exhibited the greatest intersubject variability in folded angle. This variability likely reflects substantial interindividual differences in local cortical geometry, but it may also be amplified by the relatively coarse ROI definition used here; collapsing multiple medial temporal substructures into a single ROI may pool heterogeneous folding patterns and inflate apparent orientation variability.

In the cerebellum, folded angle variability was also substantial across regions, yet its contribution to SNR difference was more modest. This attenuation is consistent with greater depth and orientation dispersion of cerebellar sources, suggesting that geometric and depth-related factors dominate detectability constraints and limit the extent to which orientation alone can drive regional SNR differences.

Importantly, while several posterior regions showed higher SNR with OPM than EEG (Fig. 4G), EEG retained higher sensitivity in other cerebellar regions. This complementary pattern indicates that OPM and EEG provide distinct but overlapping sensitivity profiles (Seedat et al., 2024), supporting their combined use to extend noninvasive evaluation beyond the cerebral cortex. This complementary EEG/MEG sensitivity profile was also observed in Samuelsson et al., 2020 (J. G. Samuelsson et al., 2020).

### 4.5 Anatomical mediation of OPM advantage

Recognizing that depth is a dominant constraint on electrophysiological detectability, we next examined whether anatomical factors mediate the relative SNR advantage of OPM-based configurations over SQUID systems in the cerebellum. By explicitly modeling sensor distance and head size, we demonstrate that anatomical factors play a key mediating role in cerebellar SNR differences between OPM and SQUID configurations.

Mediation analysis revealed that the effect of head size on SNR difference was significantly mediated by relative sensor distance in most posterior cerebellar regions (Fig. 5 F–H), whereas the anterior lobes (I–III, IV, V) and Crus I did not show significant mediation effects. These findings indicate that individuals with smaller heads—who exhibit greater asymmetry in sensor proximity between OPM and SQUID systems—derive a larger SNR benefit from cerebellar-optimized OPM layouts. This reinforces the conclusion that depth-related limitations can be partially mitigated when sensor geometry and coverage are adapted to individual anatomy, particularly for posterior cerebellar regions where OPM advantage is most evident.

These observations have important implications for clinical translation. Conventional SQUID-MEG relies on fixed helmet geometries that are optimized primarily for adult head sizes, which can further constrain sensitivity for deep sources in smaller heads. BabyMEG systems provide an important proof of concept that MEG helmets optimized for pediatric head geometries can improve MEG measurements and signal quality (Okada et al., 2016), but such systems remain specialized and are not widely available. In contrast, OPM-based systems offer a flexible alternative, enabling subject-specific sensor placement that can accommodate a broad range of head sizes while targeting regions such as the posterior fossa.

Together, these findings suggest that anatomically adaptive sensor layouts are critical for improving cerebellar detectability, and that OPM-MEG may be particularly advantageous in populations for whom fixed-helmet system are most constrained.

### 4.6 Limitations and future directions

Several limitations should be acknowledged. First, OPM results in this study were derived from simulations rather than patient recordings, and noise estimates were extrapolated from SQUID and EEG data. Although this approach was supported by a validation experiment in a healthy individual (Supplementary Fig. 1), the acquisitions were not fully equivalent: OPM and SQUID recordings were obtained in different magnetic shield room environments (single-layer versus there-layer), employed different postprocessing strategies, and involved different sensor counts. These differences limit direct quantitative generalization and motivate future work incorporating empirically measured OPM noise characteristics in clinical populations.

Second, the SNR framework relies on forward modeling, which necessarily simplifies the relationship between neural signals and noise. While this enables systematic comparison across modalities and sensor configurations, it does not capture all physiological and environmental noise contributions that may vary with tasks and recording conditions. Future studies combining subject-specific forward models with task-dependent OPM recordings will be important to refine estimates of cerebellar detectability under clinical realistic conditions.

Third, the cerebellar-optimized OPM configuration examined here was designed to enhance posterior fossa coverage but was not explicitly optimized using detailed cerebellar anatomy, functional organization, or expected source extent. More principled optimization strategies incorporating cerebellar lobular anatomy, orientation distributions, and anticipated spatial extent may yield additional improvements and represent an important direction for future cerebellar-focused OPM system design.

Finally, although we visually validated the anatomical plausibility of ARCUS-derived cerebellar segmentation against individual MRI at the lobular level, we did not directly assess geometric accuracy at the level of individual folia. Such validation would require ultra-high-resolution ground truth data not available in routine clinical imaging. Importantly, the present study was designed to evaluate the suitability of the ARCUS-derived source space for forward modeling and SNR estimation at the spatial scale relevant to M/EEG, rather than to validate folium-level surface reconstruction accuracy.

## Conclusions

Cerebellar optimized OPM configurations increased SNR particularly in posterior cerebellar regions, whereas proximity alone did not yield meaningful improvements for deep sources. Integration of individualized cerebellar modeling using ARCUS with scalp-mounted OPM arrays may help extend noninvasive electrophysiological coverage beyond the cerebral cortex. Together, these advances provide a realistic path toward improving the assessment of cerebellar involvement in epilepsy, a region that has traditionally been difficult to evaluate in clinical practice.

---

**Supple Fig. 1. Validation of brain-noise estimation for OPM simulations.** SQUID magnetometer recordings (102 channels)acquired in a three-layer magnetically shielded room and OPM recordings (14 channels) acquired in a single-layer room were obtained from the same healthy subject during somatosensory stimulation. After modality-specific processing to remove environmental magnetic interference, baseline activity from - 200 ms to -5 ms relative to stimulus onset was extracted from the evoked responses. Baseline noise was statistically compared to assess whether estimated brain-noise levels differed between SQUID magnetometers and OPM sensors.

**Supple Fig. 2. Examples of successful ARCUS-derived cerebellar segmentation.** (A, D) 24-year-old subject, 3T MRI, 1.0-mm slick thickness. (B, E) 40-year-old subject, 1.5T MRI, 1.0-mm slick thickness. (C, F) 40-year-old subject, 3T MRI, 4.5-mm slice thickness.

**Supple Fig. 3. Examples of failed ARCUS-derived cerebellar segmentation.** (A, D) 43-year-old subject, 3T MRI, 2.2-mm slice thickness. (B, E) 13-year-old subject, 3T, 0.9-mm slice thickness. (C, F) 15-year-old subject, 1.5 T, 1.2-mm slice thickness. In the first two cases, partial loss of lobules VIIb and VIIIa regions was observed (arrows). In the third case, excessively enlarged cerebellar white matter was observed.

## Abbreviations

ARCUS: Automatic Reconstruction of Cerebellar Cortex from Standard MRI Using Diffeomorphic Registration of a High-Resolution Template
BEM: boundary element model
EEG: electroencephalography
FDR: false discovery rate
MEG: magnetoencephalography
MRI: magnetic resonance imaging
OPM: optically pumped magnetoencephalography
ROI: region of interest
SD: standard deviation
SNR: signal-to-noise ratio
SQUID: Superconducting quantum interference device
SUIT: Spatially Unbiased Infratentorial Template

