## Supplementary figures and images for "Quantifying Cerebellar Signal Detectability in MEG and EEG in Epilepsy Using Anatomically Informed Source Modeling"

### supplemetary figures

Supplementary Figure 1

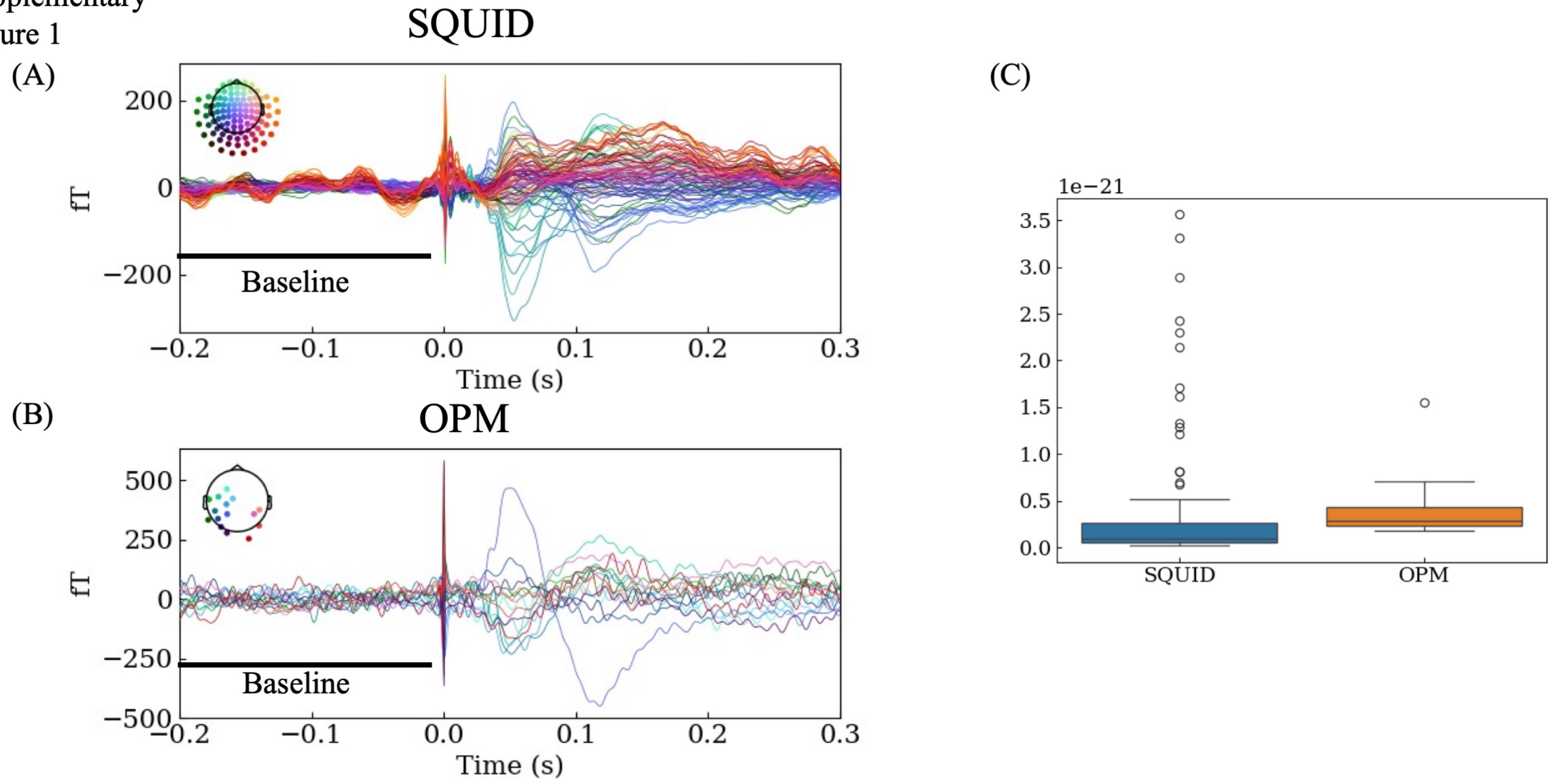

Supplementary Figure 2

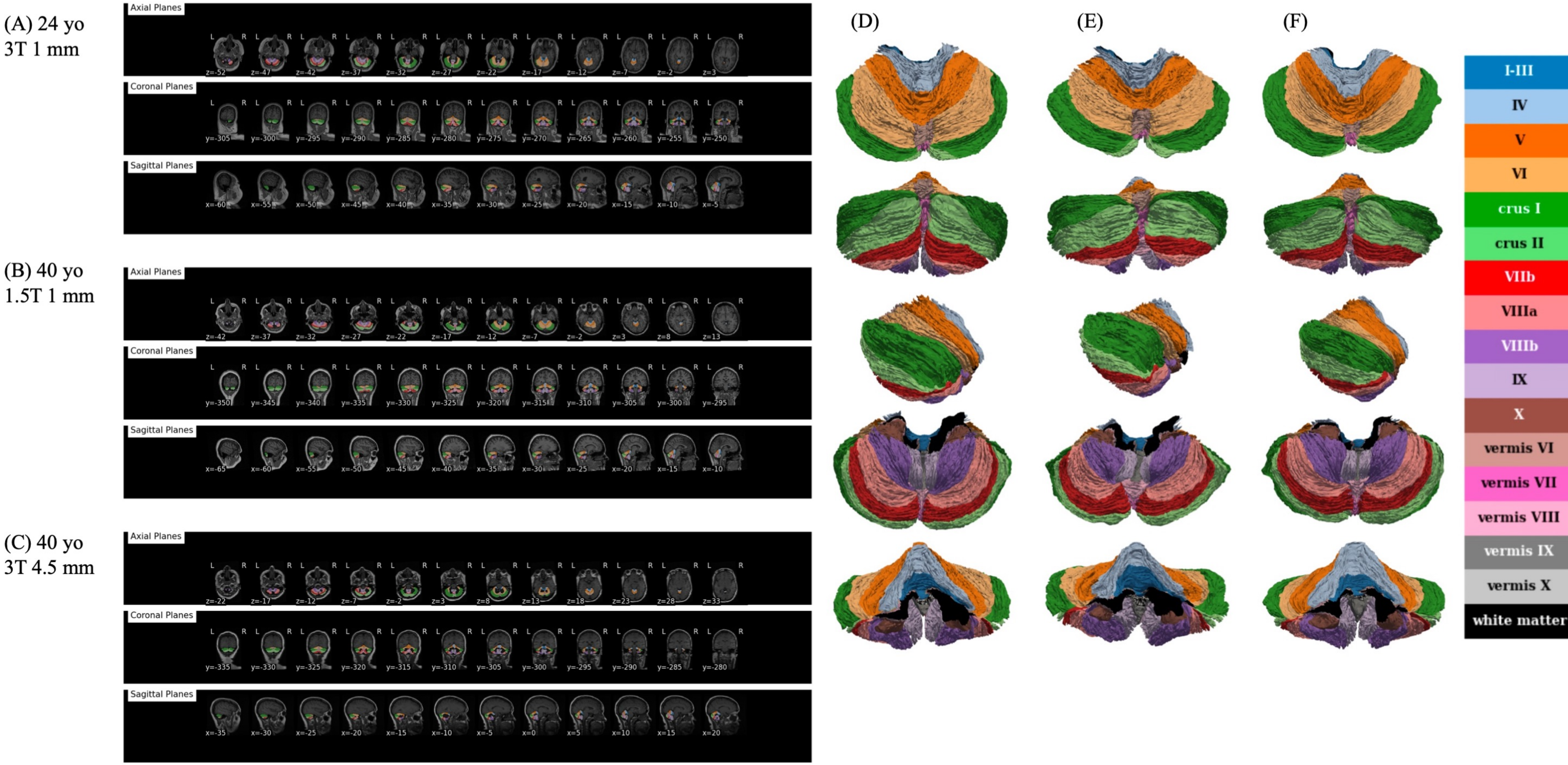

Supplementary Figure 3

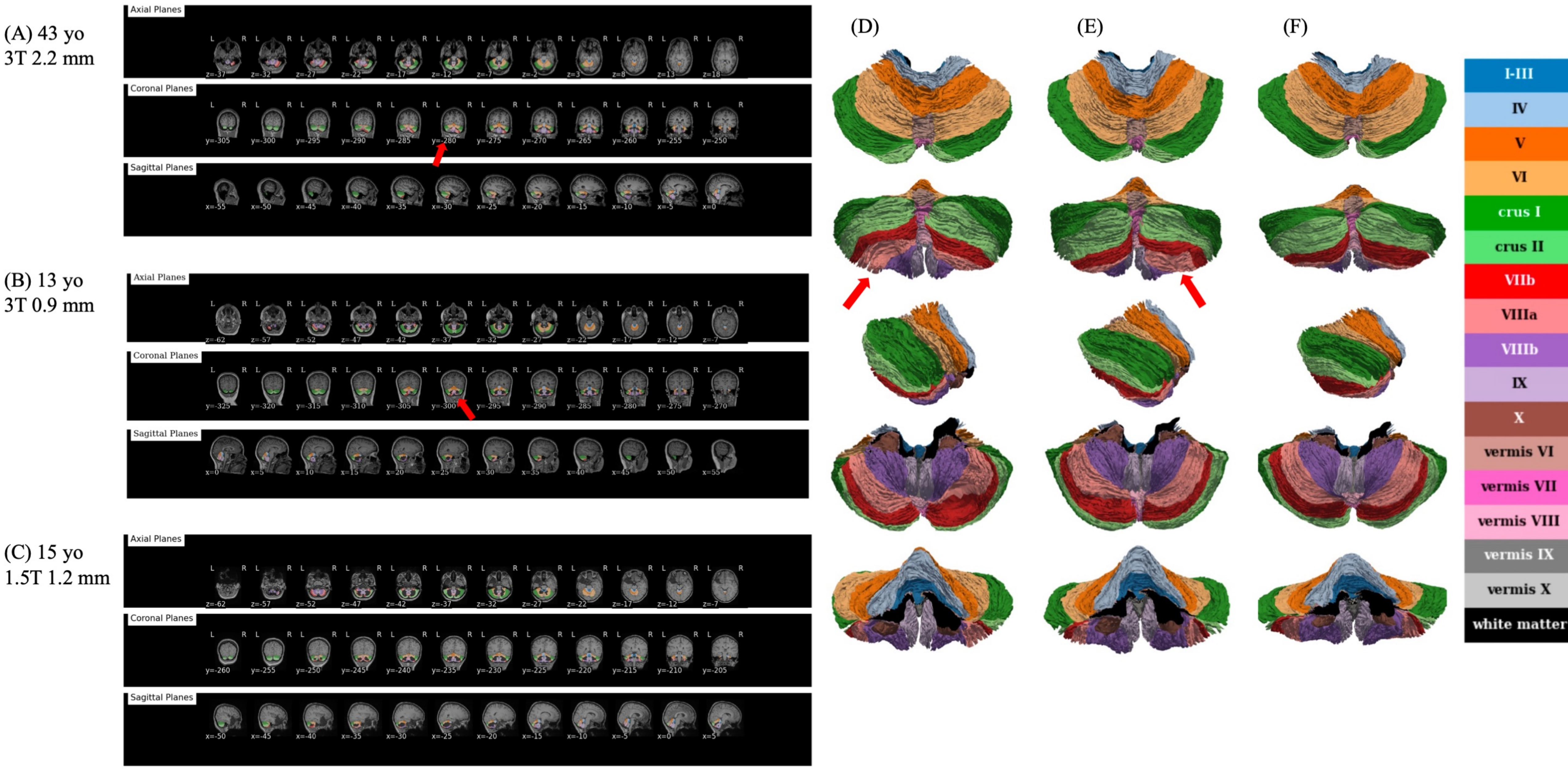
